# Single-nuclei transcriptomic analysis of the subthalamic nucleus reveals different Pitx2-positive subpopulations

**DOI:** 10.1101/677724

**Authors:** Maria Papathanou, Åsa K. Björklund, Mihaela M. Martis-Thiele, Åsa Wallén-Mackenzie

## Abstract

The subthalamic nucleus is a compartmentalized structure formed during development with each division serving a different function. A few molecular markers such as transcription factors involved in the formation and maintenance of the STN have been identified, and whilst a few more genes have been reported to be expressed in the STN, the complete transcriptomic landscape of the mature STN remains elusive. So far genes associated with the adult STN have been expressed throughout the structure. It therefore remains to establish if the functional division of this structure is attributed to a molecular diversity comprising of subpopulations. In this study a single-nuclei RNA-sequencing of genetically labelled Pitx2-Cre was conducted to identify the molecular heterogeneity of the STN. The findings revealed two subpopulations of the STN marked by the expression of Baiap3/Fxyd6 or Fgf11. Moreover additional Pitx2^+^ subpopulations with distinct signature markers that characterize neighboring structures of the STN such as the parasubthalamic nucleus, the lateral hypothalamus, the zona incerta and the mammillary nucleus were also identified.

## 1. Introduction

The subthalamic nucleus (STN) is a small excitatory structure located deep within the basal ganglia. The STN with its enriched glutamatergic projections plays a major role in motor, limbic and associative behavior (Temel et al., 2005) and is implicated in many neurological disorders, such as Parkinson’s disease (PD), supranuclear palsy and Huntington, but also psychiatric disorders such as addiction and obsessive compulsive disorder (OCD) (Dickson et al., 2010; Lange et al., 1976; Limousin et al., 2005; Pelloux and Baunez, 2013). Some of these conditions are associated with degeneration of STN neurons, whilst others are correlated with alterations in its firing pattern within the basal ganglia circuity. The STN is a common target for electrical deep brain stimulation (DBS) aimed to alleviate motor symptoms PD and essential tremor and more recently STN-DBS has been introduced for addiction and OCD. The basic mechanism underlying the therapeutic effects remains unclear. It is also uncertain why in some cases DBS is accompanied by side-effects like apathy, depression and impulsivity (Temel et al., 2006). It has been proposed that the STN can be structurally subdivided into a tripartite structure in which motor, associate and limbic divisions form different connectivity and in turn mediate distinct functions (Hamani et al., 2004), raising the possibility that the side-effects reported by patients undergoing DBS may be linked to the position of the electrode.

Previous studies have proposed that the tripartite composition of the STN arises during development, in which the neurons migrate in a spatially coordinated manner from lateral to medial throughout three sequential waves (Altman and Bayer, 1979; Philips et al., 2005). The first set of neurons migrate from the ventricular zone to the dorsolateral part of the STN and form the compartment involved in movement, whilst the associative and limbic parts consist of neurons that migrate during the second and third wave of neurogenesis, respectively. Transcription factors (TFs) are key regulators for determining the cell fate of neurons, as well their spatial and temporal acquisition, and may in turn contribute to the regional subdivision of the STN. To date a handful of TFs (FoxA1, FoxA2, FoxP1, FoxP2, Lmx1a, Lmx1b, Barhl1, Dbx1) have been shown to be expressed in subpopulations of the STN and associated with its development. However they are not exclusively linked to the STN, as some are also critical determinants of the midbrain (Gasser et al., 2016; Kee et al., 2016; Nouri and Awatramani, 2017; Skidmore et al., 2008; Zou et al., 2009). The gene encoding the TF paired-like homeodomain 2 (*Pitx2*) is expressed in the di-, mes- and rhombencephalon but serves as a key regulator for the STN. Mice lacking Pitx2 show a halted differentiation and migration of subthalamic neurons (Martin et al, 2004; Skidmore et al, 2008). Further, a Pitx2-Cre transgenic mouse line has been implemented to abrogate glutamatergic neurotransmission in the STN, which has resulted in identification of motor and reward-related features driven by the STN (Pupe et al., 2015; Schweizer et al., 2016, 2014).

To advance the understanding of molecular diversity within the STN and to identify molecular markers that could underpin the structural and functional preservation of the STN, single-nuclei RNA sequencing of the Pitx2^−positive^ population was implemented. Several subpopulations of Pitx2-positive nuclei were identified that represent, the STN and also the adjacently located parasubthalamic nucleus, the lateral hypothalamus and mammillary nucleus as well as the lateral zona incerta. Moreover we found show that in the mouse, the STN consists of two subpopulations (expressing either Baiap3/Fxyd6^+^ or Fgf11^+^) that could underlie the functional division in rodents. In summary these findings form a foundation for further structure-function analysis of the STN and neighboring structures, with the scope of improving therapeutic interventions when targeting these brain regions.

## 2. Material and Methods

### 2.1 Animal housing

Animals of both sexes were housed on a standard 12 h sleep/wake cycle (7:00 A.M. lights on, 7:00 P.M. lights off). Mice were provided with food and water *ad libitum* and housed according to Swedish legislation (Animal Welfare Act SFS 1998:56) and European Union legislation (Convention ETS 123 and Directive 2010/63/EU). All experiments were conducted with permission from the local Animal Ethical Committee.

### 2.2 Generation and genotyping of transgenic mice

Genotyping of transgenic mice was performed by PCR analysis (Supplementary Table 1). Pitx2-Cre mice (Skidmore et al., 2008) were bred with the floxed mCherryTRAP reporter line (Gt(ROSA)26Sor-mCherry-Rpl10a) (Hupe et al., 2014), generating mCherryTRAP^Pitx2-Cre^ mice.

### 2.3 Nuclear extraction and FACS sorting of mCherry-tagged nuclei

Male mCherryTRAP^Pitx2-Cre^ mice at postnatal day 28, were euthanized and the brains quickly removed and put in PBS on ice. The brain was first cut into 1mm-thick coronal section and the two subthalamic nuclei (STNs) were dissected out and snap-frozen on dry ice. Tissue was thawed and dissociated using a 1-ml dounce homogenizer (Wheaton) in ice-cold lysis buffer (0.32□M sucrose, 5□mM CaCl_2_, 3□mM MgAc, 0.1□mM Na_2_EDTA, 10□mM Tris-HCl, pH 8.0, 1□mM DTT, and 1X complete proteinase inhibitor, 0.1mM Na_2_EDTA-free (Roche)). The homogenate was slowly added over a sucrose layer (1.8□M sucrose, 3□mM MgAc, 10□mM Tris-HCl, pH 8.0, and 1□mM DTT) before a centrifugation process of 2 h 20 min at 30,000□*× g* (Beckman J-25 centrifuge with a J13.1 rotor). The supernatant was carefully aspirated and the extracted nuclei in a form of a pellet were resuspended in a nuclear storage buffer (15% sucrose, 10□mM Tris-HCl, pH 7.2, 70□mM KCl, and 2□mM MgCl_2_) containing an additional proteinase inhibitor (Complete, Roche) and an RNAse inhibitor 40 U/μL (RNAseOUT, Invitrogen). Resuspended nuclei were filtered through a 30-μm nylon cup filcon (BD Biosciences, 340625) into BSA-coated FACs tubes for sorting.

Fluorescence-Activated Cell Sorting (FACS) was performed using a BD Influx System and the BD FACS™ software (BD Biosciences) to isolate mCHERRY-tagged nuclei and single-sort into two 384 well plating containing 2.3μl of ice-cold lysis buffer with ERCC spike ins for subsequent quality control analysis. The nuclei were identified by forward- and side-scatter gating, using a 633-nm laser with a 610/20 filter. A 140-μm nozzle, a sheath pressure of 4.30□psi, an acquisition rate of up to 1200 events per second and a drop frequency of 6.05kHz were applied. Two wells of each plate were left blank and served as negative controls. The plates were quickly centrifuged after sorting and stored in −80°C until further processing.

### 2.4 Sequencing and quality control

cDNA libraries were produced and sequenced using a Smart-seq2 protocol (Picelli et al., 2013). Sequencing of the single-nuclei libraries was performed using Illumina HiSeq 2000. The expression values were computed as reads per kilobase of gene model and million mappable reads (RPKMs) to normalize for varying sequencing depths across sequenced nuclei and the gene lengths. The expression values of merged RefSeq and Ensemble gene annotations were computed as described in (Ramsköld et al., 2009), using uniquely aligned reads and correcting for the uniquely alignable positions using MULTo (Storvall et al., 2013). The reads were mapped and aligned to mouse genome (mm10) with the STAR aligner (Dobin et al., 2013) using 2-pass alignment to have improved performance of de novo splice junction reads, filtered for only uniquely mapping reads. Only exons were included in the analysis. Parameters for quality control included uniquely mapping reads (<80%), ERCC detection (< mean −2 standard deviations), ERCC ratio (>0.1) fraction of reads mapping to exons (<10%), detected with RPKM>1 (>3000 and < mean + 2 standard deviations) and a maximum correlation to another nuclei (< 0.3).

### 2.5 Bioinformatic analysis

The obtained RPKM data was analyzed using the Seurat package 2.3.4 in R (Butler et al., 2018) to identify subpopulations followed by MAST (Model-based Analysis of Single Cell Transcriptomics) (Finak et al., 2015) to find differentially expressed genes (significantly up- or downregulated) between the individual clusters. Differentially expressed genes were visualized using the T-embedded stochastic neighbor embedding (t-SNE), which is a non-linear dimensionality reduction method and violin plots or dotplots. A minimum of 3 nuclei and minimum of 200 genes was a preliminary criterion for the Seurat object. For the analysis the Seurat package was run twice following the removal of one cluster that did not meet the neuronal criteria. For the two runs variable gene selection was set for the x-low, x-high and y cut-off at 0.8, 10 and 1 for the first run and 0.6, 10 and 1 for the second run, respectively. Clustering was run with the default SNN clustering at resolution of 0.8. The number of Principal components (PCs) used for the analysis was based on a Jackstraw analysis for the first 12 PCs with a cut-off of 0.001. For the first run PCs 1-7 and 12 were used, whilst for the second Seurat only the first 6 PCs were included.

### 2.6 Histological analysis

#### 2.6.1 Immunohistochemistry

Deeply anaesthetized mice were transcardially perfused with body-temperature phosphate-buffered saline (PBS) followed by ice-cold 4% formaldehyde. Brains were dissected and post-fixed overnight. The brains were then cryo-protected with 30% sucrose and cut using a cryostat at 30µm slice thickness. Free-floating sections were processed for immunofluorescence according to standard protocols (Primary Antibodies: guinea-pig Calbindin D28K (CALB1) 1:350 #214-004, Synaptic Systems; rabbit Calretinin (CALB2) 1:500 #NBP1-88221, Novus Biologicals; rabbit Adcyap1 1:54 #OACD01441 Nordic Biosite, rabbit Stard5 1:175 #NBP1-92448, Novus Biological. Secondary antibodies: Donkey Anti-Rabbit Alexa Fluor 488 1:500; Donkey-Anti-Guinea-Pig Alexa Fluor 488 1:500; DAPI 1:5000). mCHERRY was detected by the endogenous fluorescence without any additional use of an antibody. Images were captured using a NanoZoomer *20.2-HT.0* and processed using the Ndp2.view software (Hamammatsu).

## 3. Results

### Single cell isolation and transcriptomic analysis of glutamatergic Pitx2-Cre positive cells of the STN

To resolve the molecular composition of the STN at the genome-wide gene expression level, a transgenic mouse-line carrying the enzyme Cre-recombinase under control of the Pitx2-promoter (Skidmore et al., 2008) was crossed with the reporter line floxed mCherryTRAP reporter line (Gt(ROSA)26Sor-mCherry-Rpl10a) (Hupe et al., 2014) to generate mCherryTRAP^Pitx2-Cre^ mice. Expression of mCherry was strongly detected in the STN but also in the posterior, lateral and ventromedial hypothalamus (Fig. 1A). The expression resembled the reported data of *Pitx2* mRNA in the Allen Brain Atlas (Probe RP_071018_03_B04). Accumulation of the tagged L10a protein in nucleoli allows for purification of nuclei via FACS after nuclear preparation (Heiman et al., 2008). Both STNs from 4 mice were dissected and the extracted nuclei sorted in two 384 well plates, each containing nuclei from two animals. Libraries from single nuclei were generated using Smart-seq 2 protocol (Picelli et al., 2013) for full-length RNA sequencing (snRNAseq) at a depth of 200-250 million reads and with 50bp per single read (Fig. 1B & C). About 80% of the reads showed unique alignments to the mouse genome with 20% of the reads aligning within exons. Following a strict quality control (QC), 138 nuclei did not meet the criteria in two of the six QC parameters and were excluded from further analysis (Supplementary Figure 1A). Consistent between the two experimental plates, the median number of counts was 354578 and median number of detected genes per nuclei was on 5384 for the remaining 630 nuclei (Supplementary Figure 1B & C). Out of the 18697 genes found across the 630 nuclei, 1776 were selected as variable genes (Supplementary Figure 1D).

**Figure 1.**
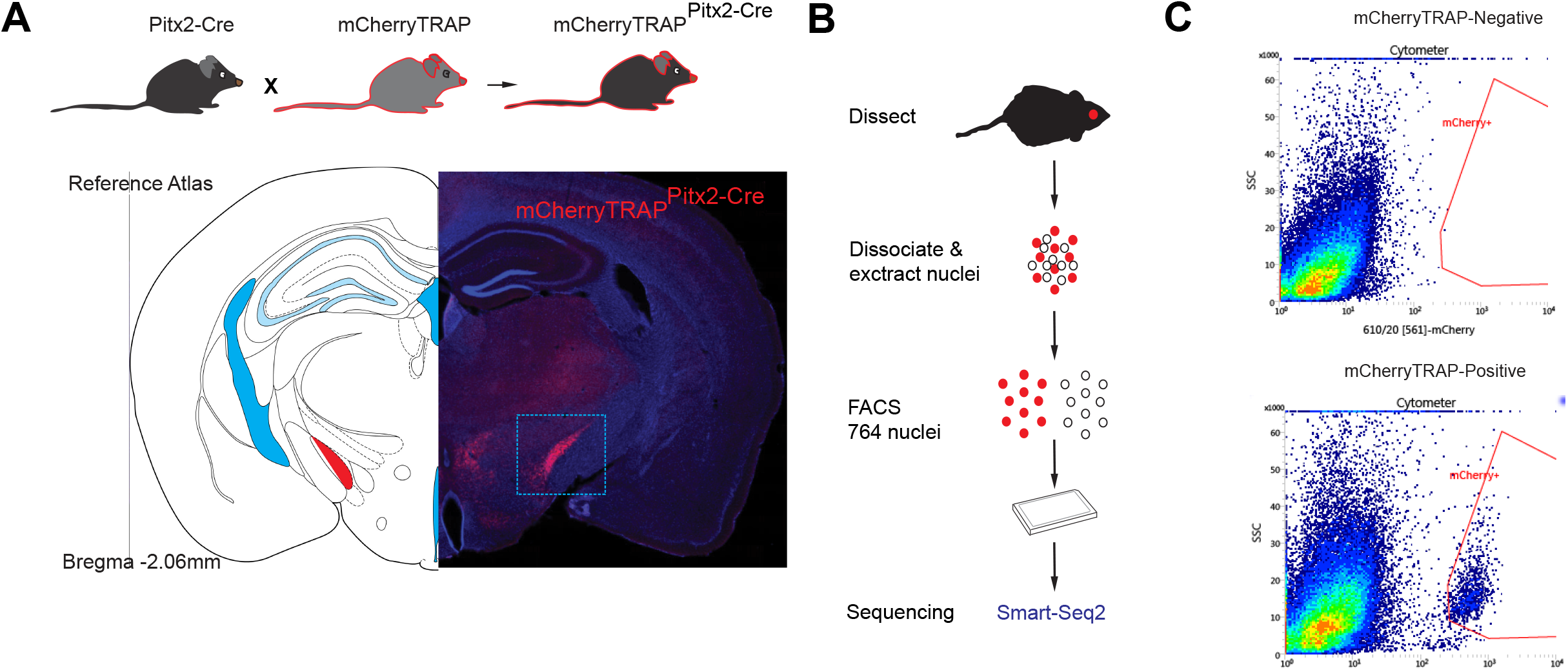
(**A)**: Schematic illustration of breeding strategy of Pitx2-Cre mice and the mCherryTRAP reporter to generate mCherryTRAP^Pitx2-Cre^ mice. mCherry expression in nucleoli of transgenic mice and region of dissected tissue shown in inset. **(B)** Schematic illustration of experimental procedure from dissection to sequencing. **(C)** FACS plot of mCherry negative and positive population.

The Seurat package was employed in order to identify subpopulations and unique markers within each cluster from our snRNAseq dataset. The 630 samples were clustered into 6 groups with 185,174, 114, 101, 45 and 11 nuclei in clusters 0 to 5, respectively (Fig. 2A). The number of detected genes was similar across the clusters, with the exception of cluster 5, which contained nuclei with high RPKM values (Fig. 2B). On the first instance, *Pitx2, Slc17a6* and *Slc32a1* were visualized in the obtained t-SNE-plot to verify that the dataset is of the correct population. As expected, all clusters showed to be of glutamatergic lineage and Pitx2-positive and sparsely expressed the vesicular inhibitory amino acid transporter (Fig. 2C-E). Similarly other known TFs associated with the STN, such as *Lmx1a, Foxa1, Foxp1, Foxp2* and *Barhl1* were also expressed in most clusters, although surprisingly *Lmx1b* was barely detected. In the dark-blue cluster (cluster 4) only *Foxp2* was expressed but none of the other markers, suggesting possibly a Pitx2-positive population other than the STN (Supplementary Figure. 2A-F).

**Figure 2.**
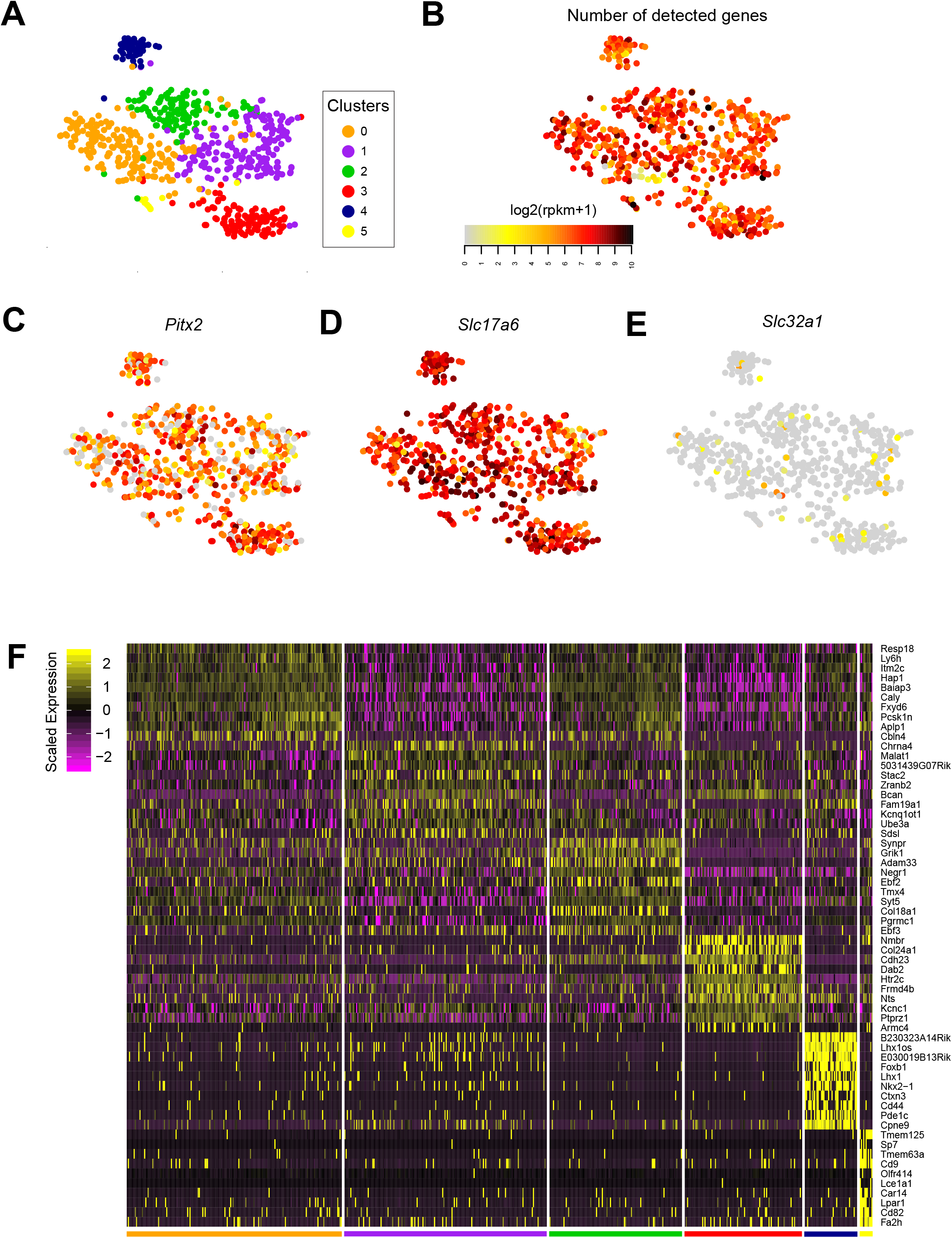
**(A)** t-SNE embedding with 6 clusters (orange, purple, green, red, dark blue and yellow) **(B)** Number of detected genes per nuclei (expression levels ranging from light grey (low) to black (high) **(C-E)** Expression levels of *Pitx2, Slc17a6* and *Slc32a1* among the 6 identified clusters as visualized in the t-SNE plot **(D)** Heatmap showing the top ten genes considered unique for and characterizing each cluster

Of the 6 clusters, cluster 3, 4 and 5 clearly separated from the rest, whilst clusters 0-2 appeared to be more closely related (Fig. 2D). This was evident in the violin plots although uniquely identified markers stood out for each group (Supplementary Figure 2G-L). Importantly, since the signature markers for cluster 5 were associated with the oligodendrocyte lineage (*Tmem125, Tmem63a, Cd9, Car14, Lpar1*), this population was excluded from further analysis and the filtered dataset subjected to a second round of unsupervised clustering with Seurat.

### Identification of Pitx2-positive subpopulations of the STN and neighboring structures

In the second run 1946 variable genes were detected and the remaining 619 nuclei grouped again in 6 groups (Fig. 3A). Among the top 10 genes characterizing each cluster, *Baiap3, Hap1, Irx3, Cacna2d3* and *Ctxn1* were in the orange cluster, *Chrn4a, Stac2, Bcan, Fam19a1* and *Sdsl* in the purple cluster, *Synpr, Grik1, Adam33, Ebf2* and Tac1 in green cluster, *Nmbr, Col24a1, Cdh23, Htr2c* and *Frmd4b* in the red cluster and *Cartpt, Foxb1, Lhx1, Nkx2-1* and *Ctxn3* in the dark blue cluster pointing to subthalamic, hypothalamic as well as mammillary nuclei (Fig. 2E & F). These five clusters matched those obtained from the first Seurat run as rated from the top 10 genes distinguishing each cluster and were therefore similarly color coded in the t-SNE plot and the heatmap (Fig. 3A & B). The 10 genes characterizing the additional cluster were *Rtn, Tmem130, Fxyd6, Resp18, Caly, Pcsk1n, Gaa, Tmem59l, Itm2c* and *Ndfip1*. Given that this subpopulation showed close resemblance to the green cluster, this group was color-coded light-green (Fig. 3B). When looking at the number of nuclei there was a reduction in the size of the orange cluster (now 92 nuclei) compared to the remaining ones, suggesting that the new light-green cluster with 104 nuclei may be spatially similar to the orange as well as the green cluster.

**Figure 3.**
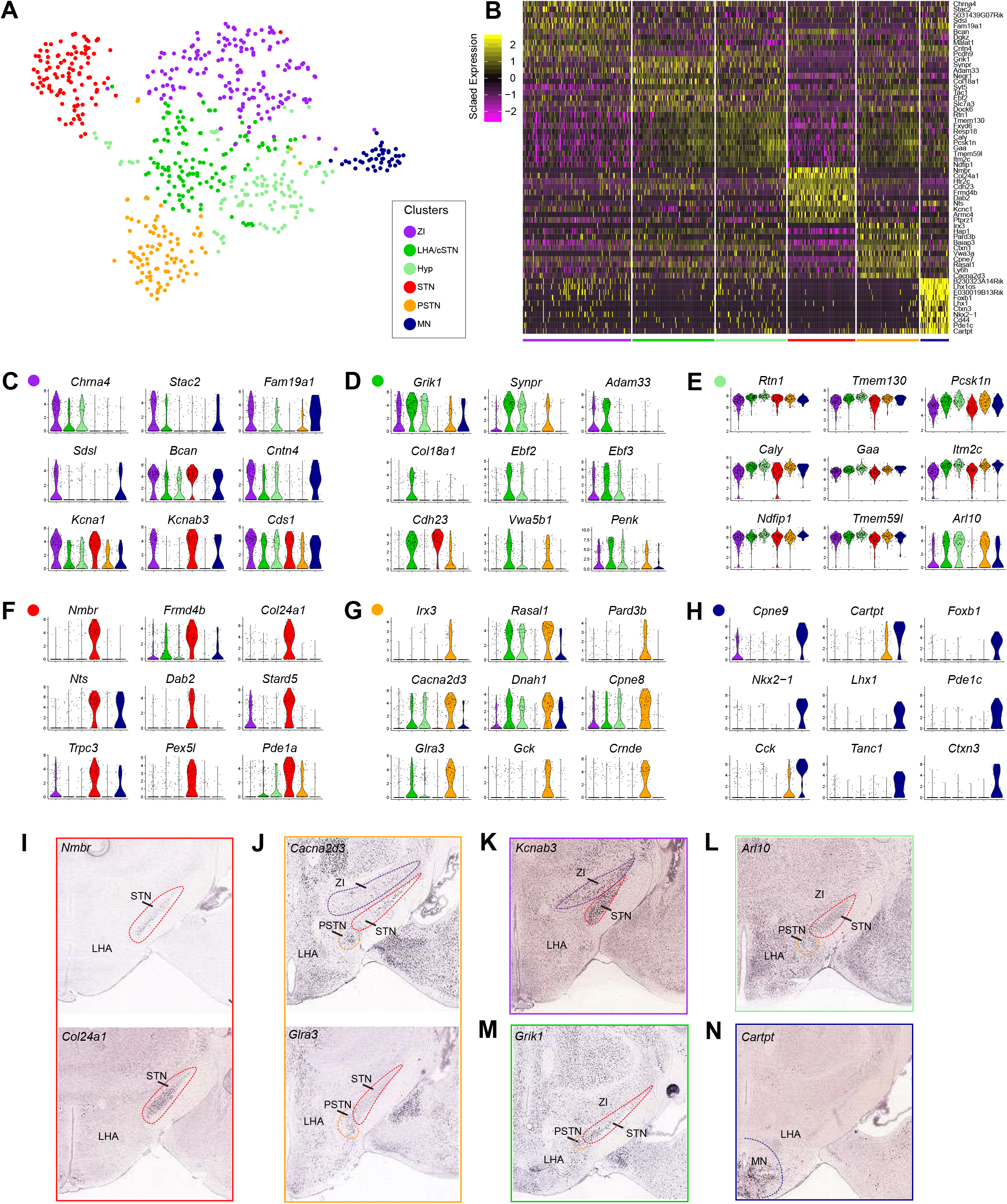
**(A)** t-SNE embedding of 6 clusters (purple, green, light green, red, orange, dark blue). **(B)** Heatmap showing top ten genes considered unique for or characterizing each cluster **(C-H)** Violin plots of six discriminatory markers of each populations **(I-L)** In situ images taken from the Allen Brain Atlas for candidate genes (representing the different clusters: *Nmbr*: RP_070116_02_A04; *Col24a1*: RP_050915_04_C05, *Cacna2d3:* RP_071204_04_A05, *Glra3:* RP_051213_01_B04; *Kcnab3*: RP_050927_01_A02; *Grik1*: RP_060518_02_E09; Arl10: RP_060220_04_C02; *Cartpt*: RP_051017_03_B03. Abbreviations: Hyp, Hypothalamus; LHA: Lateral hypothalamic area; MN: Mammillary nucleus PSTN: Parasubthalamic nucleus; cSTN, caudal Subthalamic nucleus; STN: Subthalamic nucleus; ZI: Zona incerta

In an attempt to find out what exactly these Pitx2-Cre positive populations represent, a differential expression analysis between the groups was run with MAST and subsequently top candidates validated with the Allen Brain Atlas. The number of significantly upregulated genes detected when comparing one cluster versus the rest with a p adjusted value of <0.01 were 19 (purple), 25 (green), 50 (light green), 141 (red), 69 (orange) and 94 (dark-blue) for the different clusters (Supplementary Figure 3), whilst the downregulated genes were 87, 15, 30, 149, 16 and 69 for the 6 clusters, respectively. Interestingly, when comparing the significantly upregulated genes between the two green and the orange clusters, *Cdh23* was the only gene differing between the two green clusters, whilst *Grik1, Ebf2, Dmb, Adam33, Plk5, Ntm, Ebf2* and *Ephb1* (green cluster) and *Rtn1, Syngr1, Ntm* and *Serinc1* (light-green) were upregulated genes compared to the orange cluster. Conversely, *Nxph1, Vwa3a, Irx3, Pcsk4, Adamts2, Pcdh17, Cacna2d3, Cpne8, Ddc, Nnat* (green), *Dbn1* (light-green) and *Cpne7, Stxbp2* (in both) were downregulated compared to the orange cluster.

When comparing these genes with the Allen Brain Atlas and with previous datasets looking at hypothalamic populations (Chen et al., 2017; Mickelsen et al., 2019) it appeared that the light-green population marked general hypothalamic neurons (Fig. 3E, L and Supplementary Figure 3), whilst the green cluster consisted of markers *(Synpr, Adam33, Col18a1, Calb2, Ebf3* and *Penk*) of the lateral hypothalamic area (LHA). However in the recent study by Mickelsen et al., (2019) *Ebf3, Synpr* and *Pitx2* made up three distinct LHA subpopulations, suggesting that in our case the green cluster might characterize a mixture of the very caudal STN (cSTN) and the neighboring Pitx2-positive nuclei of the LHA (Fig. 3D, M & Supplementary Figure 3). The orange cluster with markers like *Glra3, Irx3, Cacna2d3, Pard3b* and *Gck* seems to represent the parasubthalamic nucleus (PSTN) that were mostly expressed in this cluster albeit in lower percentage of cells, whilst *Hap1* and *Baiap3* were highly expressed in most cells but also present in the other hypothalamic clusters (Fig. 3G, J & Supplementary Figure 3).

Genes associated with the purple cluster were *Chrna4, Stac2, Fam19a1, Bcan, Kcna1* and *Kcnab3*, the former of which has previously been reported to be expressed in the STN during development (Agulhon et al., 1998). When looking at the Allen Brain Atlas most of these were detected in the lateral Zona incerta (ZI). However some of these genes shared similar expression levels to the red cluster (Fig. 3C), which appears to characterize the STN in particular the anterior STN, raising the possibility that this population may also be dorsolateral subgroup of the STN (Fig. 3C & K and Supplementary Figure 3).

Ample of genes such as *Nmbr, Col24a1, Dab2, Trpc3, Pex5l* were almost exclusively expressed in the subthalamic population (red cluster). *Frmd4b, Stard5, Robo2, Kcnc1, Neto2, Cdh23* and *Htr2c* were highly expressed in almost all neurons of the STN and with partial expression in the other clusters, whilst *Nts* was predominantly expressed in this and the final dark-blue cluster (Fig. 3 F, I & Supplementary Figure 3). This latter cluster was primarily characterized by TFs such as *Foxb1, Nkx-2-1, Lhx1* and markers like *Cartpt, Cck* and *Tanc1*, and whilst many have been reported in several regions of the medial hypothalamus *Foxb1* is distinct for the mammillary nucleus (MN), (Zhao et al., 2008), thus suggesting that this is the Pitx2-positive population of the MN (pre and retromammillary).

### *Baiap3*/*Fxyd6* and *Stard5/Fgf11* define two subpopulations of the STN

When visualizing genes of the general hypothalamic population like *Rtn1, Tmem130, Pcsk1n, Fxyd6, Gaa* and *Caly* onto the t-SNE (Fig. 3E), it was evident that despite them being detected in all clusters that there was a gradient expression from high to low along the axis of tSNE-2, particularly with *Fxyd6* and *Caly* (Supplementary Figure 4). Indeed when analyzing more of the differentially expressed genes, markers *Baiap3, Fxyd6, Syt5, Gpx3, Sprint2, Camkv* and *Ctxn1* were highly expressed in the MN, the hypothalamus, the LHA/cSTN, the PSTN clusters and importantly, within the STN, only in the nuclei located in the bottom part of the STN cluster of the t-SNE (Fig. 4B_1-7_). Given the anatomical distribution of these brain regions, suggests that *Baiap3, Fxyd6* and *Syt5* may not only distinguish the STN along the rostrocaudal axis but possibly also along the mediolateral axis. Indeed in accordance to the *in situ* data from the Allen Brain *Baiap3* and *Fxyd6* were expressed in the very medial part of most the STN and in the cSTN (Fig. 4C, D), raising the possibility that this molecular subpopulation may also underlie the functional division reported within the STN.

**Figure 4:**
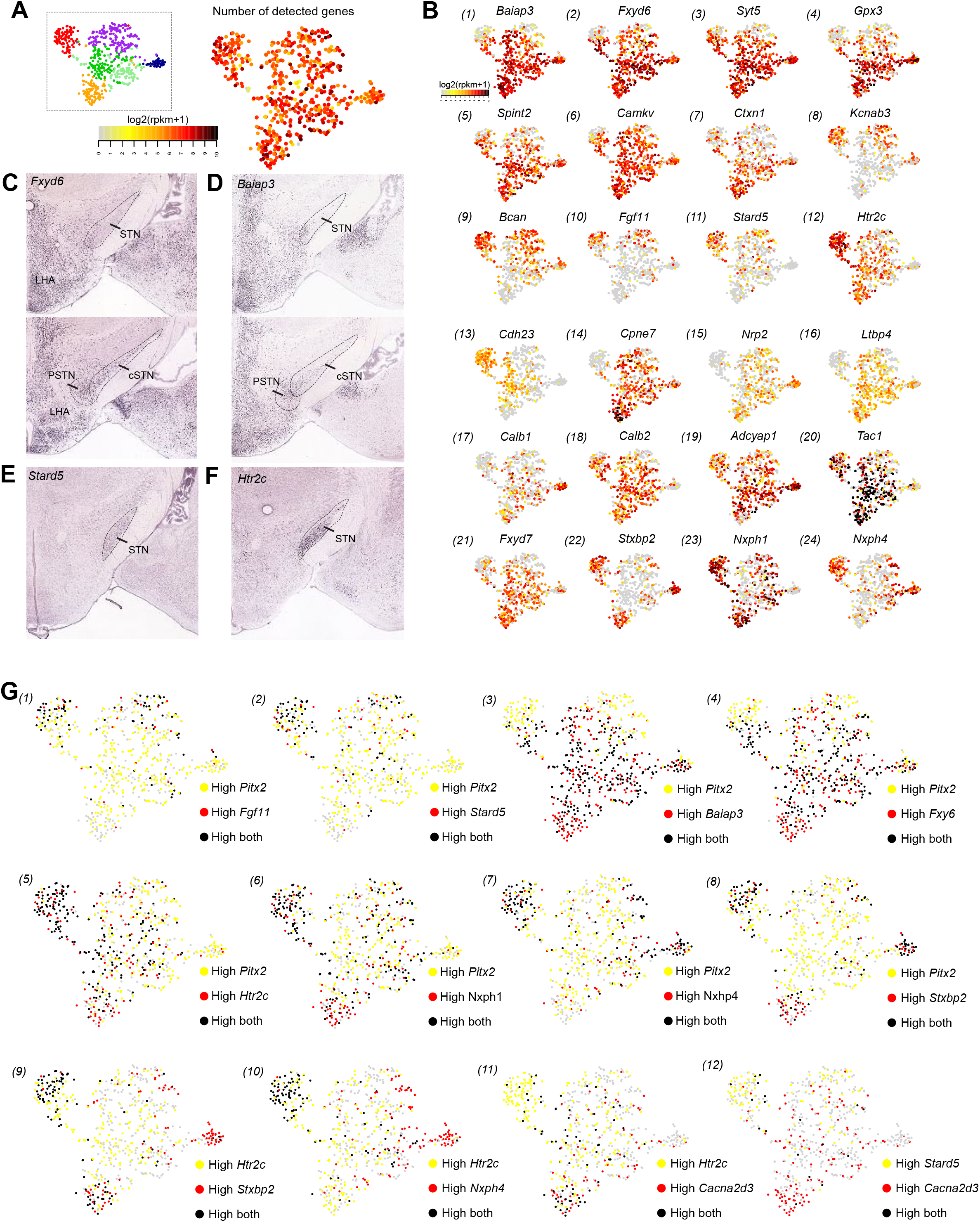
**(A)** Number of detected genes detected per nuclei (expression levels ranging from light grey (low) to black (high) **(B**_**1-24**_) tSNE-plots of selected differentially expressed genes **(C-F)** In situ images taken from the Allen Brain Atlas for *Fxyd6* (RP_051017_01_E10), *Baiap3* (RP_060315_03_C11), *Stard5* (RP_050610_01_A03) and *Htr2c* (RP_050825_02_B06) **(G**_**1-12**_) Overlay of differentially expressed genes with Pitx2 and other combinations showing combinatorial expression unique for subpopulations.

Conversely *Kcnab3, Bcan, Fgf11* and *Stard5* were predominantly expressed in the lateral ZI cluster and most of the STN cluster but sparsely expressed in the other clusters including the PSTN (Fig. 4B_8-11_, E). A finding that was further validated when looking at the protein level in which STARD5 was expressed in the STN but not in the PSTN and LHA (Fig. 5D-D’’’). Importantly *Fgf11* was expressed in nuclei residing in the exact opposite end of the STN cluster compared to those expressing *Baiap3* and *Fxyd6*, thereby making these three markers possible candidates for STN subpopulations (Baiap3^+^/Fxyd6^+^/Fgf11^−^ and Baiap3^−^/Fxyd6^−^/Fgf11^+^) (Fig. 4B_8-11_, E).

**Figure 5:**
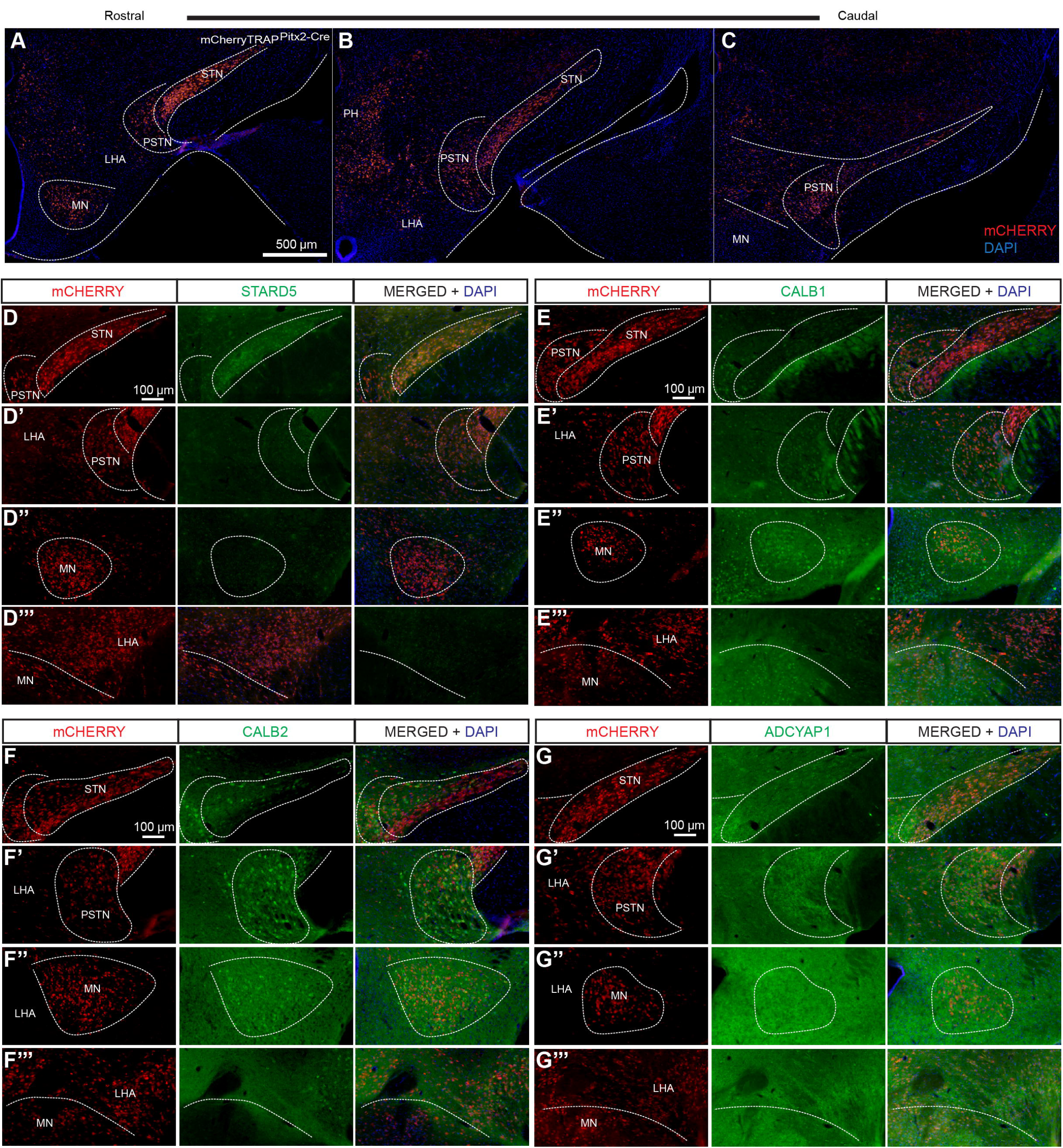
**(A-C)** Expression of mCHERRY in mCherryTRAP-Pitx2-Cre along the rostrocaudal axis **(D-D’’’)** Expression of mCHERRY and STARD5 in Pitx2-Cre positive subpopulation **(E-E’’’)** Expression of mCHERRY and CALB1 in Pitx2-Cre positive subpopulation **(F-F’’’)** Expression of mCHERRY and CALB2 in Pitx2-Cre positive subpopulation **(G-G’’’)** Expression of mCHERRY and ADCYAP1 in Pitx2-Cre positive subpopulation Abbreviations: LHA: Lateral hypothalamic area; MN: Mammillary nucleus PSTN Parasubthalamic nucleis; cSTN, caudal Subthalamic nucleus; STN: Subthalamic nucleus; PH, Posterior hypothalamus

Similarly to the gradient expression between the tSNE-2 axis, there was also a difference along the tSNE-1 axis, with *Cdh23* and *Htr2c* strongly expressed in the STN, LHA/cSTN and PSTN but absent from the more caudal Pitx2-population of the MN (Fig. 4B_12-13_, F). In contrast markers like *Cpne7, Nrp2, Ltbp4* and *Calb1* were higher in the PSTN, hypothalamus and MN clusters but very lowly expressed in the STN, whilst *Adcyap1, Calb2, Tac1, Fxyd7* were broadly expressed in the PSTN and hypothalamic populations including LHA/cSTN but appeared in salt and pepper manner within the STN (Fig. 4B_14-21_ & Fig. 5E-G). Importantly, we were able to identify markers like *Stxbp2* and *Nxph1* that were expressed in the STN and PSTN clusters but lowly detected or absent from the LHA/cSTN and *Nxph4* expressed solely in STN but not in the PSTN, thereby adding a further molecular distinction between the STN and the PSTN (Fig. 4B_21-24_). In summary instead of solely using *Pitx2* as a candidate marker using combinatorial expression of genes such as *Fgf11, Stard5, Baiap3, Fxyd6, Htr2c, Nxph1, Nxph4, Stxbp2* and *Cacna2d3*, will enable further differences between the neuronal populations that make up the lZI, the MN, the STN, the PSTN and the LHA/cSTN (Fig. 4G_1-12_, Fig. 6).

**Figure 6:**
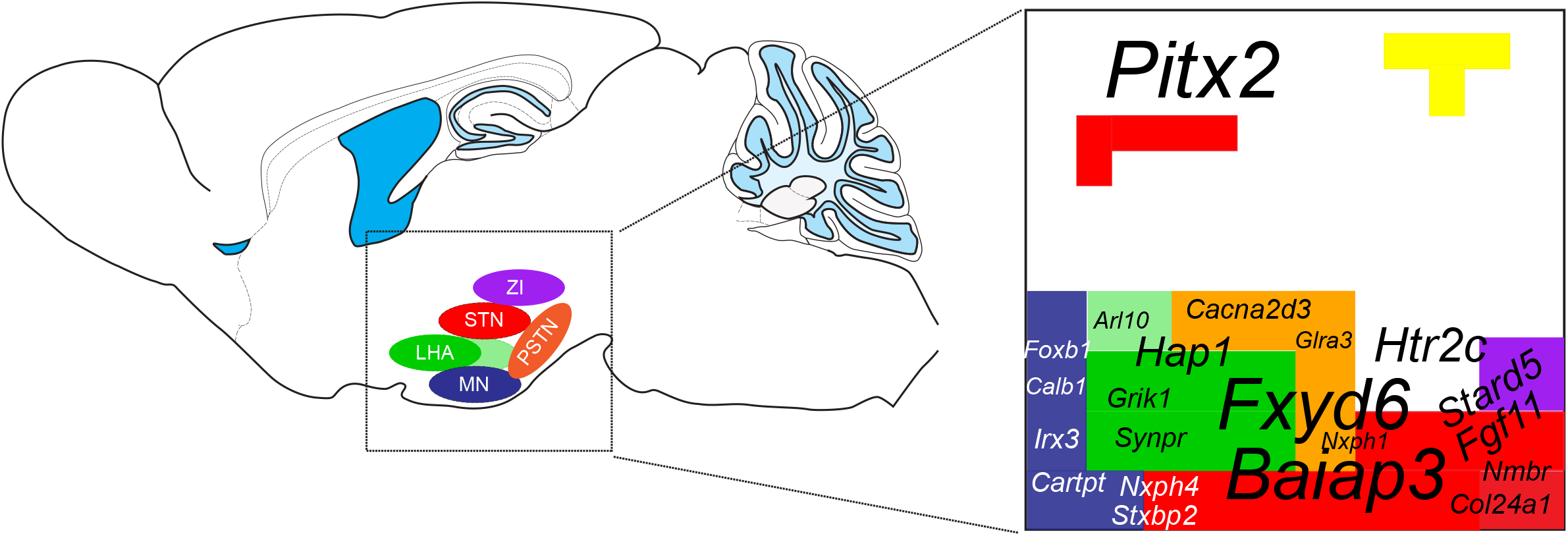
Sagittal section of a mouse brain depicting the identified Pitx2-Cre positive subpopulations with a summary of differentially expressed genes for the each of these clusters, which are similarly color-coded. Markers detected in more than one cluster are indicated by overlapping among the different colored shapes. Abbreviations: LHA: Lateral hypothalamic area; MN: Mammillary nucleus PSTN Parasubthalamic nucleis; STN: Subthalamic nucleus

## 4. Discussion

Recent studies have explored the diversity of the hypothalamus (medial and lateral) by means of single cell RNA-seq (Chen et al., 2017; Mickelsen et al., 2019; Romanov et al., 2016). However, the identification of neuronal subtypes characterizing the STN, located in proximity to the hypothalamus, is however mostly unexplored, which adds a certain degree of limitation when studying the STN in the context of neural circuits, behavior and disease. In this study, a snRNA-sequencing approach of genetically labelled Pitx2-Cre population was employed to unravel the molecular landscape and heterogeneity of STN, with the scope of identifying whether the structural and functional division is a result of different neuronal subpopulations. Six Pitx2-positive subpopulations were identified representing the MN, general hypothalamic populations, LHA/cSTN, PSTN, STN and ZI based on discriminate markers such as *Foxb1, Fxyd6, Synpr, Cacna2d3, Col24a1* and *Chrna4* that were upregulated in the individual clusters and their anatomical expression pattern evaluated from the Allen Brain Atlas. Importantly markers *Baiap3* and *Fxyd6* appeared to characterize one subpopulation of the STN cluster, whilst *Fgf11* marked the second one. In addition compared to *Pitx2, Stxbp2, Nxph1* and *Nxph4* seemed to distinguish between the STN, PSTN, LHA/cSTN and MM with *Stxbp2* expressed in all but the LHA cluster, *Nxph1* in the STN and PSTN and *Nxph4* in STN and MN. Finally, in this study a comprehensive gene list (*Nmbr, Frmd4b, Dab2, Stard5, Pex5l, Galnt14, Trpc3)* was found to be mostly expressed in the STN cluster and together with *Htr2c, Nts, Nxph4*, and *Cdh23* can be used to study the STN in a more selective manner than *Pitx2*, which is expressed throughout the STN, but also in the neighboring structures.

*Pitx2* has been shown to be expressed in the dorsal midbrain, the pre- and retromammilllary nuclei, various regions of the hypothalamus and the STN and to be a key regulator for the development of the STN but also the superior colliculus (Martin et al., 2004). However, whilst the development of the STN is most commonly thought to arise from the posterodorsal hypothalamic neuroepithelium or the closely related mammillary nucleus, some report the STN to develop from the ventrolateral thalamic neuroepithelium and consider the STN to be an intermediate zone of the ventral thalamus (Marchand, 1987; Philips et al., 2005). Interestingly, the different theories regarding the origin of the STN were also reflected in the results of the present study. Whilst most of the identified clusters are of hypothalamic nature or reside close to the hypothalamus such as the PSTN and the MN, the purple cluster that represents the ZI is marked by genes (eg. *Kcnab3, Kcna1, Stard5, Bcan*) that are shared between this and the red STN cluster, could potentially explain a more thalamic nature. Moreover, it seems that the t-SNE1 axis depicts the rostrocaudal migratory path with the red more anterior STN cluster and the MN cluster residing on the two opposite ends of the axis. This is also supported by the expression of *Pitx2* and *Calb1* and their co-localization detected in the MN. The t-SNE2 axis on the other hand probably reflects the dorsoventral or mediolateral pattern with the ZI and PSTN clusters in the two ends of the axis. mCherry^Pitx2-Cre^ neurons were detected in the posterior and lateral hypothalamus, the MN, the PSTN and the STN, which is in accordance to the Allen Brain Atlas but also to previous studies addressing the expression of Pitx2 in development (Kee et al., 2016; Skidmore et al., 2008). However the current dataset provides further molecular markers that can distinguish between these brain regions beyond the TFs (eg. Lmx1a, Foxa1, Foxp1, Foxp2, Barhl1) identified in the aforementioned two studies. Of note, whilst the aforementioned TFs and Irx3 denoted the subthalamic lineage in the study by Kee *et al*. 2016, in the present study Irx3 was uniquely found in the PSTN cluster, although the possibility of sharing common progenitor lineage cannot be excluded. Moreover in this study the genes identified for the various clusters were associated with ion channels, transmembrane proteins, cell surface protein mediating, neuropeptides and neurotransmitter receptors among others with only the MN cluster showing a higher number of TFs. This finding is however not too surprising given that the mice used in this study were 28 days old, in which genes linked with cellular machinery rather than the differentiation process may be more essential.

*Baiap3* and *Fxyd6* were highly expressed in most clusters, similarly to previously broadly expressed markers *Adcyap1* and *Calb2* (Chen et al., 2017; Romanov et al., 2019). However the former were computationally distinctly expressed in only half of the red STN cluster. According to the Allen Brain Atlas, *Baiap3* and *Fxyd6* was expressed in the more medial part of the anterior STN. In the rodent this ventromedial part of the STN with its reciprocal connections to the ventral pallidum has been linked to associative and limbic functions (Tan et al., 2006). Given the anatomical expression of *Baiap3* and *Fxyd6*, suggests that these genes may play a role in more limbic functions. So far *Baiap3* has been shown to be significantly downregulated in the STN of mice exposed to 28 days of MPTP toxicity that could relate changes in STN firing with the transcript levels of *Baiap3* (Lauridsen et al., 2011) but its role in behavior remains unknown. Whilst it is still unclear whether the functional division with the different afferents and efferents of the STN are associated to gene expression, the current finding of the two subpopulations within the red STN cluster, marked by *Baiap3*^*+*^/*Fxyd6*^*+*^ *or Fgf11*^*+*^ makes these genes prime candidates for further studies.

High frequency stimulation of the STN often results in adverse effects like depression which is believed to be linked with associated with serotonin depletion through inhibition of the midbrain serotonergic neurons (Temel et al., 2007). Interestingly, in the current study the second most upregulated gene of the red STN cluster compared to the rest is the serotonin receptor (Htr2c). The rather high expression of *Htr2c* in the STN is in accordance to the Allen Brain Atlas and scRNA-seq dataset by (Wallace et al., 2017). As such this particular marker could be used further to explore the interplay of the STN and the serotonin system and possibly shed light in the adverse events reported with STN-DBS.

So far, little is known about the PSTN. It resides medially to the STN yet it is remains unclear how distinct it is from the STN and the LHA. One study reported the PSTN to influence hindbrain components of the central parasympathetic control network and to play a role feeding behavior and cardiovascular regulation, whilst differing from the LHA by a distinct population expressing β-preprotachykinin (Goto and Swanson, 2004). The findings obtained from the present study show that markers *Cpne7, Hap1, Gpx3* and *Fxyd7* that are highly expressed in the PSTN are also found in the LHA/cSTN cluster and general hypothalamic cluster, *Stxbp2* and *Nxph1* found in both STN and PSTN but not LHA/cSTN whilst markers like *Ctxn1, Cacna2d3, Rasal1, Glra3* and *Gck* predominantly found in the PSTN cluster. As such all these markers in addition to *Nxph4* that is found in the STN but not the other two clusters could be used in future studies to elucidate further the structural and functional role of the PSTN and possibly delineate the similarities or differences between the PSTN and the STN or LHA.

To conclude whilst many studies so far have picked up the Pitx2-positive subpopulation of the LHA or have looked at the EP in conjunction with the STN, this is the first study that fully addressed the heterogeneity of the STN and Pitx2-Cre positive subpopulations. The identification of these differentially expressed genes and unique markers for the STN, PSTN, LHA/cSTN and MM can now be explored further in the context of anatomy and by using their promoters as regulatory elements in Cre-driver lines for functional experiments implementing Cre-Lox transgenics to achieve selectivity, e.g. conditional genetics, optogenetics and chemogenetics.

## Acknowledgments

The authors thank Professior James Martin, Baylor College of Medicine (Houston, TX, USA) for sharing the Pitx2-Cre mice and Jan Stenman (former Ludwig Institute for Cancer Research, Stockholm Branch) for the mCherryTRAP reporter mice. Konstantinos Toskas and Dr Erik Södersten are thanked on advice on nuclear isolation carried out at the Holmberg Lab (Ludwig Institute for Cancer Research, Stockholm Branch and CMB, Karolinska Institutet). The authors thank Jaromir Mikes for technical assistance at the FACS Facility at Science for Life Laboratory. The single-cell transcriptome data was generated at the Eukaryotic Single-cell Genomics facility at Science for Life Laboratory in Stockholm, Sweden. The computations were performed on resources provided by SNIC through Uppsala Multidisciplinary Center for Advanced Computational Science (UPPMAX) under Project b2017161 and SNIC 2017/7-252.

## Author contributions

M.P: Performed experiments and bioinformatics analysis, analyzed data, prepared figures and wrote manuscript; Å.K.B: Supervised bioinformatics analysis; M.M: Performed bioinfomatics analysis; Å.W.M: Supervised study and wrote manuscript. All authors read the paper and provided input.

---

**Supplementary Table 1.**
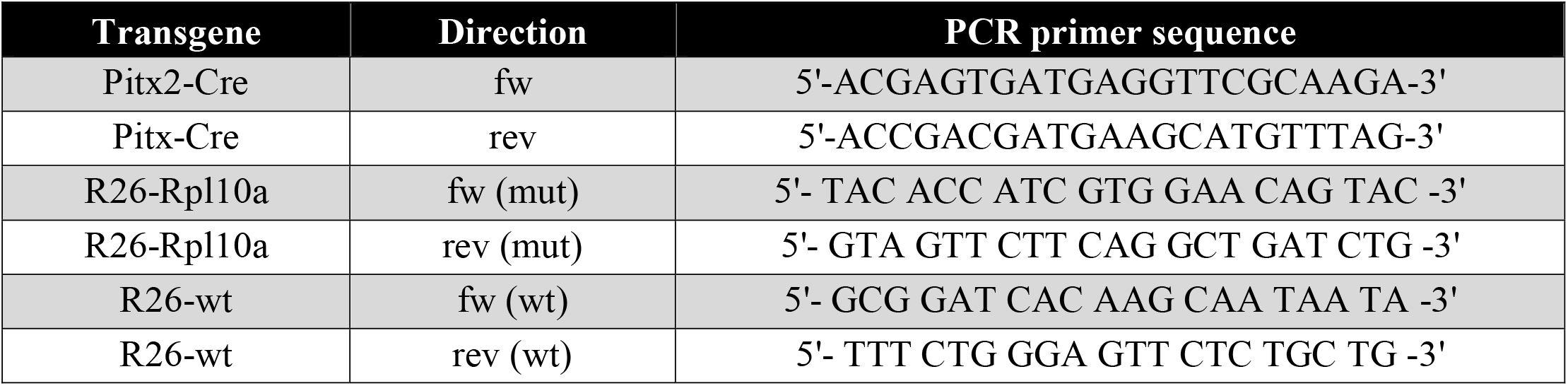
PCR primer sequences used for genotyping of transgenic mice employed in the study.

**Supplementary Figure 1 related to Figure 2: (A)** Quality control of the two experimental plates with histograms of percent uniquely mapping (Unique), Spike-in detection (ERCC), Spike-in ratio (ERCC-ratio), percent mapped to exons (Exons), detected genes with RPKM>1 (RPKM) and highest correlation to another cell (Max correlation). Nuclei which failed cutoffs are colored in red **(B)** Violin plots showing the number of detected genes of each nuclei across the two experiment plates according to Seurat unsupervised clustering **(C)** Violin plots showing the average number of transcripts across the two experiment plates **(D)** Identification of highly variable genes. The dispersion of the genes (variance/log-average expression) is plotted against the average expression (normalized log-expression, on x-axis). The minimum threshold to be considered highly variable was set at a dispersion of 1 on the y-axis.

**Supplementary Figure 2 related to Figure 2: (A-F)** Expression levels of knowns transcription factors associated with the STN visualized in the t-SNE plots. **(G-L)** Violin plots of 5 of the top 10 identified discriminatory markers of each clusters

**Supplementary Figure 3 related to Figure 3 and 4: (A-F)** Dotplots of up to 25 significantly upregulated genes of each cluster vs the rest using MAST R package

**Supplementary Figure 4 related to Figure 4:** tSNE-plots of **(A)** *Rtn1*, **(B)** *Tmem130* **(C)** *Pcsk1n*, **(D)** *Caly* **(E)** *Gaa* **(F)** *Fxyd6* showing gradient of expression along the t-SNE-2 axis

