## Supplementary figures and images for "Single-nuclei transcriptomic analysis of the subthalamic nucleus reveals different Pitx2-positive subpopulations"

### Supplementary Figure 1

Supplementary Figure 1 related to Figure 2

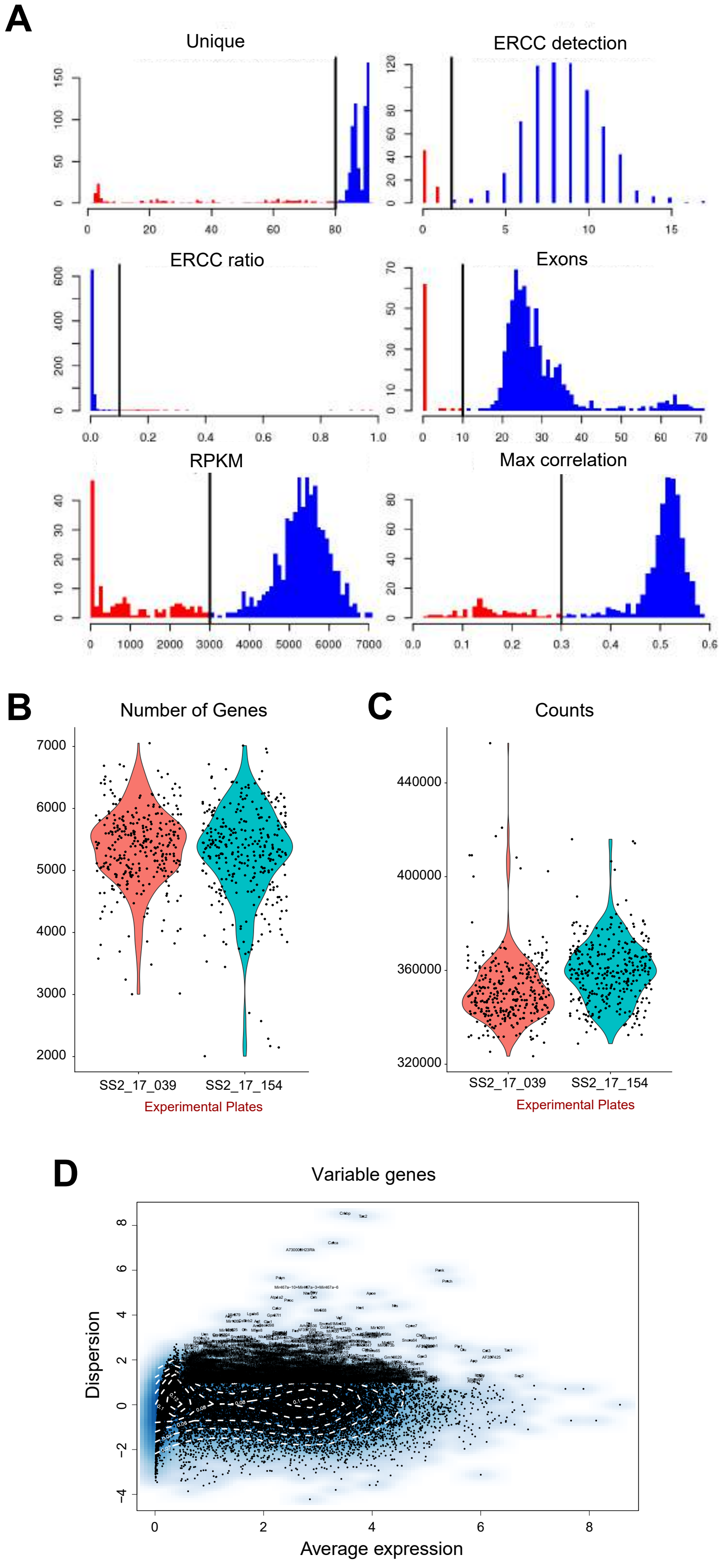

### Supplementary Figure 2

Supplementary Figure 2 related to Figure 2

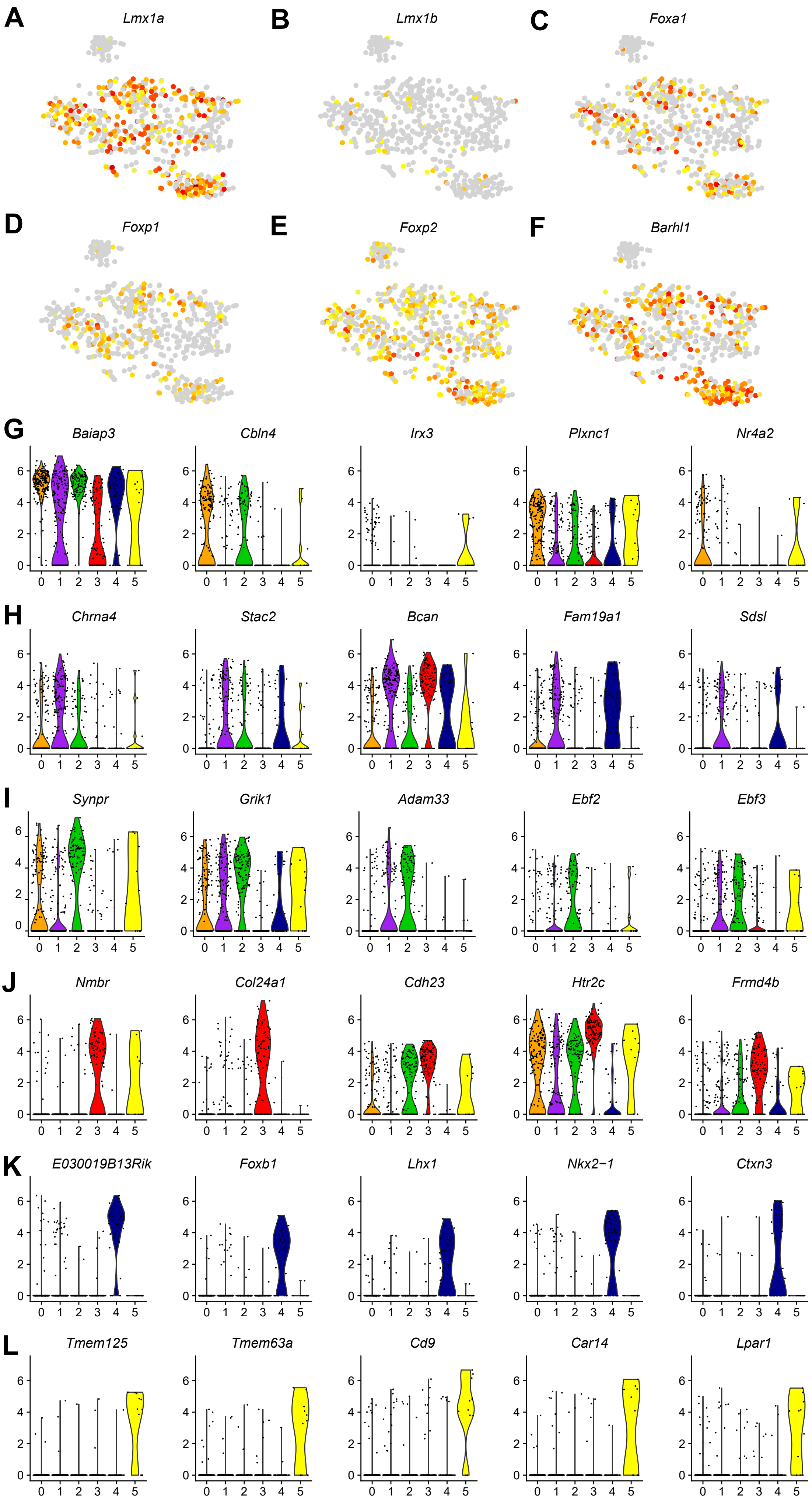

### Supplementary Figure 3

Supplementary Figure 3 related to Figure 3 and 4

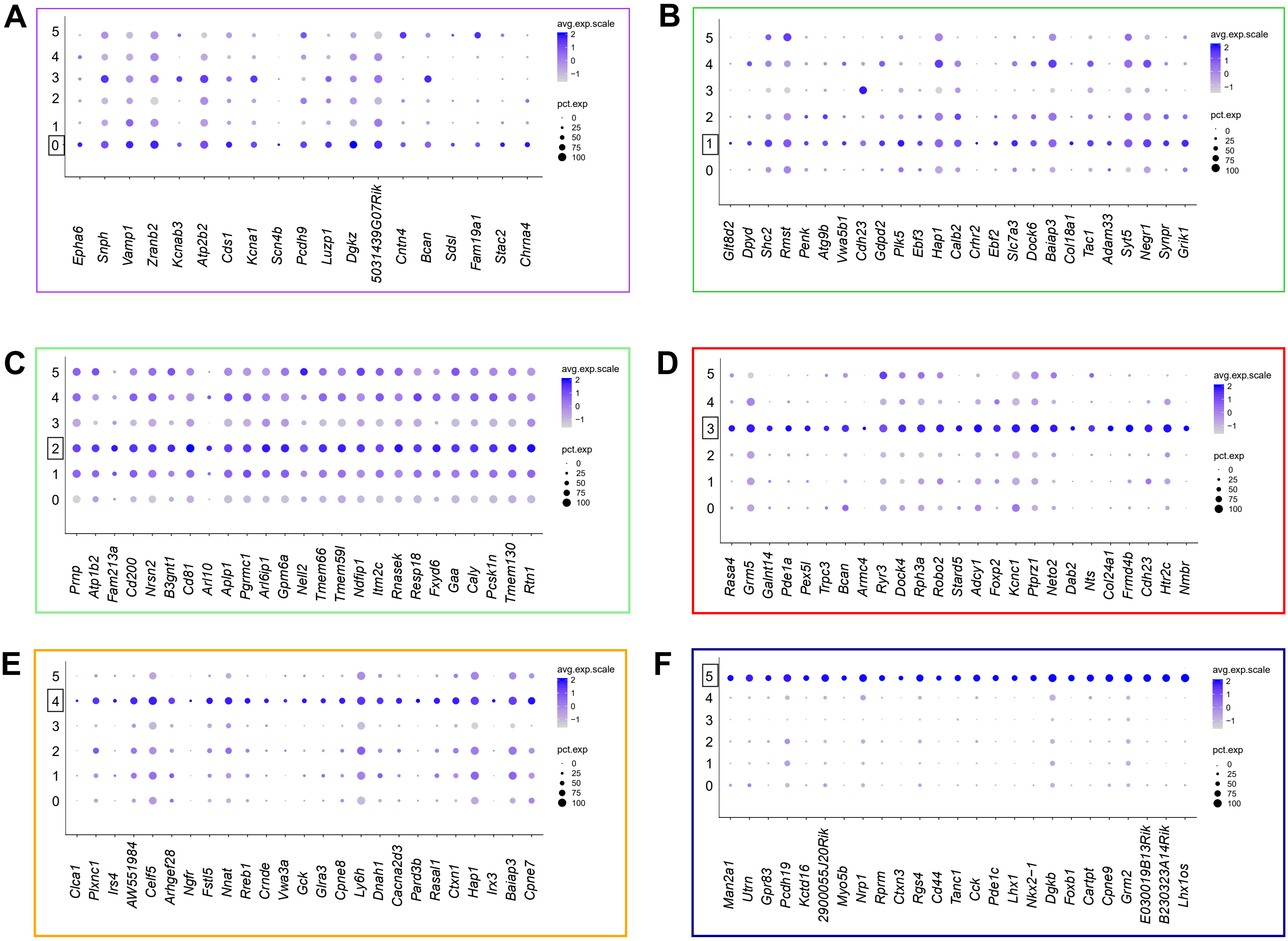

### Supplementary Figure 4

Supplementary Figure 4 related to Figure 3 and 4

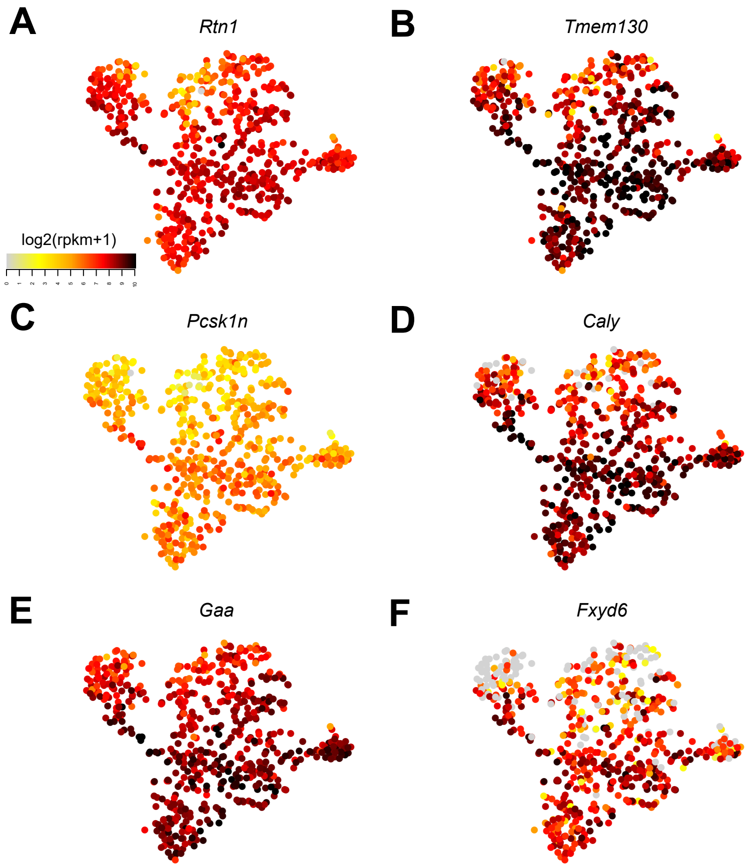
